# Urbanization and green corridors influence reproductive success and pollinators of common milkweed

**DOI:** 10.1101/2022.03.11.483986

**Authors:** Sophie Breitbart, Albert Tomchyshyn, Helene Wagner, Marc Johnson

## Abstract

Urbanization exerts many pressures on species, yet little is known about how these pressures impact species interactions. Studies of urban plant-pollinator systems provide mounting evidence that urbanization impairs pollinator movement in fragmented urban landscapes, yet the consequences for pollinator-mediated plant reproduction remains unclear. In non-urban areas, habitat corridors can facilitate the movement of organisms including pollinators, but whether these corridors facilitate plant-pollinator interactions in urban areas remains understudied. To examine how urban environments and green corridors influence plant-pollinator interactions, we measured reproductive success in the native plant common milkweed (*Asclepias syriaca*), and the community structure of its pollinators, for two years along two urban-rural transects in the Greater Toronto Area, Canada, one of which followed a green corridor. We found that urbanization decreased male fitness (i.e., pollen removal), increased fruit set (i.e., mean no. of follicles per inflorescence), and inconsistently affected female fitness (i.e., no. of follicles) in *A. syriaca*. Urbanization simultaneously decreased pollinator abundance but increased pollinator richness. Proximity to a green corridor inconsistently affected male fitness but increased reproductive effort (i.e., no. of inflorescences) in *A. syriaca*, while pollinator diversity and richness was lower in corridors. Notably, there were no consistent relationships between pollinator community structure and reproductive success in *A. syriaca* in both the presence, and absence, of a green corridor. These results demonstrate the complexity with which urbanization, green corridors, and pollinator communities can shape the reproductive investment and fitness of native plant populations.

## Introduction

Urbanization causes profound ecological change driven by intense interacting anthropogenic stressors. Urban ecosystems undergo many abiotic ecological changes including increased pollution, reduced pervious surface area, and elevated temperatures (Grimm et al. 2015; McDonnell & MacGregor-Fors 2016). These changes occur in concert with biotic changes such as altered species composition, increased habitat fragmentation, and a higher prevalence of introduced species (Faeth et al. 2005; McKinney 2006; El-Sabaawi 2018; Miles et al. 2019). Although recent advances have greatly clarified how urban environments influence the ecology of many species, we still have little understanding of how urbanization affects species interactions and their consequences for individuals, populations or communities (Shochat et al. 2006; Turrini et al. 2016). Perturbations to species interactions such as pollination can influence individuals by reducing survival rates and fecundity, which can influence population growth rates and potentially the structure and stability of communities (Heithaus 1974; Brown et al. 2001; Barraclough 2015). Thus, understanding how urban environments restructure species interactions is essential for sustaining complex networks of communities and conserving the ecosystem services they provide.

Plant-pollinator systems are an excellent model for understanding how urbanization influences species interactions. Plants and their pollinators include a wide breadth of functional traits that can be influenced by urbanization in various ways (Harrison & Winfree 2015; Aronson et al. 2016; Wenzel et al. 2019; Prendergast et al. 2022). Several aspects of pollinator ecology, including species abundance and distribution, habitat availability, body size, and foraging habits, are prone to disruption by elements of the urban landscape (Harrison & Winfree 2015; Wenzel et al. 2019; Theodorou et al. 2020a; Proesmans et al. 2021; Prendergast et al. 2022). In particular, widespread habitat loss and fragmentation, increased pollution, proliferation of introduced species, and warmer temperatures associated with urbanization may lead to changes to plant-pollinator interactions (Harrison & Winfree 2015; Wenzel et al. 2019). For instance, phenological mismatch now occurs in some urban areas because of shifts to earlier flowering in plants but not earlier pollinator flight activity (Fisogni et al. 2020). As another example, urban air pollution significantly reduced pollinator abundance and flower visits in experimental populations of *Brassica nigra* (Ryalls et al. 2022). Thus, the variation of pollinator and plant responses to urbanization makes plant-pollinator systems ideal for testing hypotheses about how urbanization affects species interactions more generally.

Changes in urban plant-pollinator interactions can have important consequences for the reproductive success of insect-pollinated plants. Preliminary evidence suggests urban plants and pollinators often experience parallel, positive or negative, fitness impacts depending on the response of specific pollinators and plants to urban environmental change (Rivkin et al. 2020). For example, most studies indicate that urbanization negatively affects plant reproductive success as a result of altered pollinator availability and/or habitat fragmentation (Cheptou & Avendaño 2006; Pellissier et al. 2012; Geslin et al. 2013; Ushimaru et al. 2014; Hermansen et al. 2017; Oliveira et al. 2019). However, the generalist-pollinated *Centaurea jacea*, in its native range, experienced higher reproductive success in isolated urban parks than urban semi-natural sites, illustrating how some plants can respond positively to urbanization (Van Rossum 2010). It is generally accepted that improving ecological conditions for urban pollinators is likely to bolster reproductive success in insect-pollinated plants and facilitate urban plant-pollinator interactions.

Habitat corridors can provide important refugia and connectivity for maintaining species interactions and reproductive success. In the case of plant-pollinator interactions, the fragmented urban landscape can disrupt pollination and negatively impact plant fitness (Cheptou & Avendaño, 2006; Aguilar et al. 2006; Andrieu et al. 2009). However, it may be possible to increase functional connectivity in urban environments with green infrastructure known as ecological corridors: generally long, narrow strips of land that can facilitate dispersal among otherwise isolated patches (Aars & Ims 1999; Haddad et al. 2003). Previous research on invertebrates has largely focused on how ecological corridors affect species diversity (Angold et al. 2006; Lynch 2019; Twerd et al. 2021). We are unaware of any studies that examine how these corridors impact urban plant-pollinator interactions or pollinator communities specifically.

Here, we tested the hypothesis that urban environments and green corridors influence pollinator communities and consequently plant reproduction by studying natural populations of a native plant, common milkweed (*Asclepias syriaca*), and its pollinators along an urbanization gradient. We asked four specific questions: 1) How does urbanization influence a plant ‘s investment in reproduction and reproductive success? 2) How do green corridors in urban and suburban areas affect plant reproduction? 3) Does the pollinator community structure of *A. syriaca* change along an urbanization gradient? 4) How do green corridors in urban and suburban areas impact pollinators? Answering these questions will provide insight into how urbanization and green corridors together shape pollinator communities and natural plant population fitness in urban environments.

## Methods

### Study System

Common milkweed, *Asclepias syriaca*, is an herbaceous perennial plant native to eastern North America. *Asclepias syriaca* is an obligate outcrosser, reproducing sexually via hermaphroditic flowers, and it also spreads vegetatively via rhizomes to produce clonal ramets (Wilbur 1976; Wyatt & Broyles 1994). Plants often grow in old fields and along roadsides, where they occur as discrete patches of one ramet to several thousand ramets (Wilbur 1976). Inflorescences consist of a cluster of ca. 50 flowers arranged in an umbel (Thompson et al. 2017), and each flower contains five pollinaria: sets of two adjoining pollen sacs (pollinia; Bhowmik & Bandeen 1976). Pollination occurs after pollinaria are removed by insects (e.g., Hymenoptera, Lepidoptera; Wilbur 1976), and then deposited into the stigmatic slit of another milkweed flower from a different plant (Willson & Price 1977; Wyatt & Broyles 1994). Cross-fertilization yields follicles (i.e., fruits) that grow until they dehisce and release wind-dispersed seeds, whereas self-fertilized follicles typically abort before reaching 4–6” (Bhowmik & Bandeen 1976; Willson & Price 1977). *Asclepias syriaca* is an apt model organism for investigating urban plant-pollinator interactions for several reasons: plants are common along urban-rural gradients, fruit production (an estimate of female fitness) depends on insect pollinators, and it is possible to quantify pollen removal as an estimate of male fitness by counting the number of discrete pollen sacs removed from flowers.

### Field Sampling

To understand the effects of urbanization and green corridors on plant reproduction, we sampled plants at 80 sites along an urban-rural gradient in the Greater Toronto Area, Ontario, Canada (Figure 1). This 67 km transect extended from downtown Toronto southwest towards Hamilton, Ontario, and was split into three subtransects: two parallel urban subtransects and one rural subtransect, all roughly equal in length. The urban subtransects extended west from the urban center and were separated by ca. 5 km, with the northern subtransect (“Urban: Non-Corridor”) traversing the urban matrix. The southern subtransect (“Urban: Corridor”) followed a mostly continuous vegetated corridor ca. 50–350 m wide running along utility rights-of-way and railroad tracks with sampling sites within or nearby the corridor. The rural subtransect (“Rural”) extended from the urban subtransects ‘ western ends. Sampling sites (henceforth “populations”) contained discrete patches of *A. syriaca* and were chosen from a variety of habitats such as farmland buffers, parks, residential gardens, and natural areas based on the availability of plants, with the constraint that populations were spaced >500 m apart to increase their independence (Pleasants 1991).

**Fig. 1.**
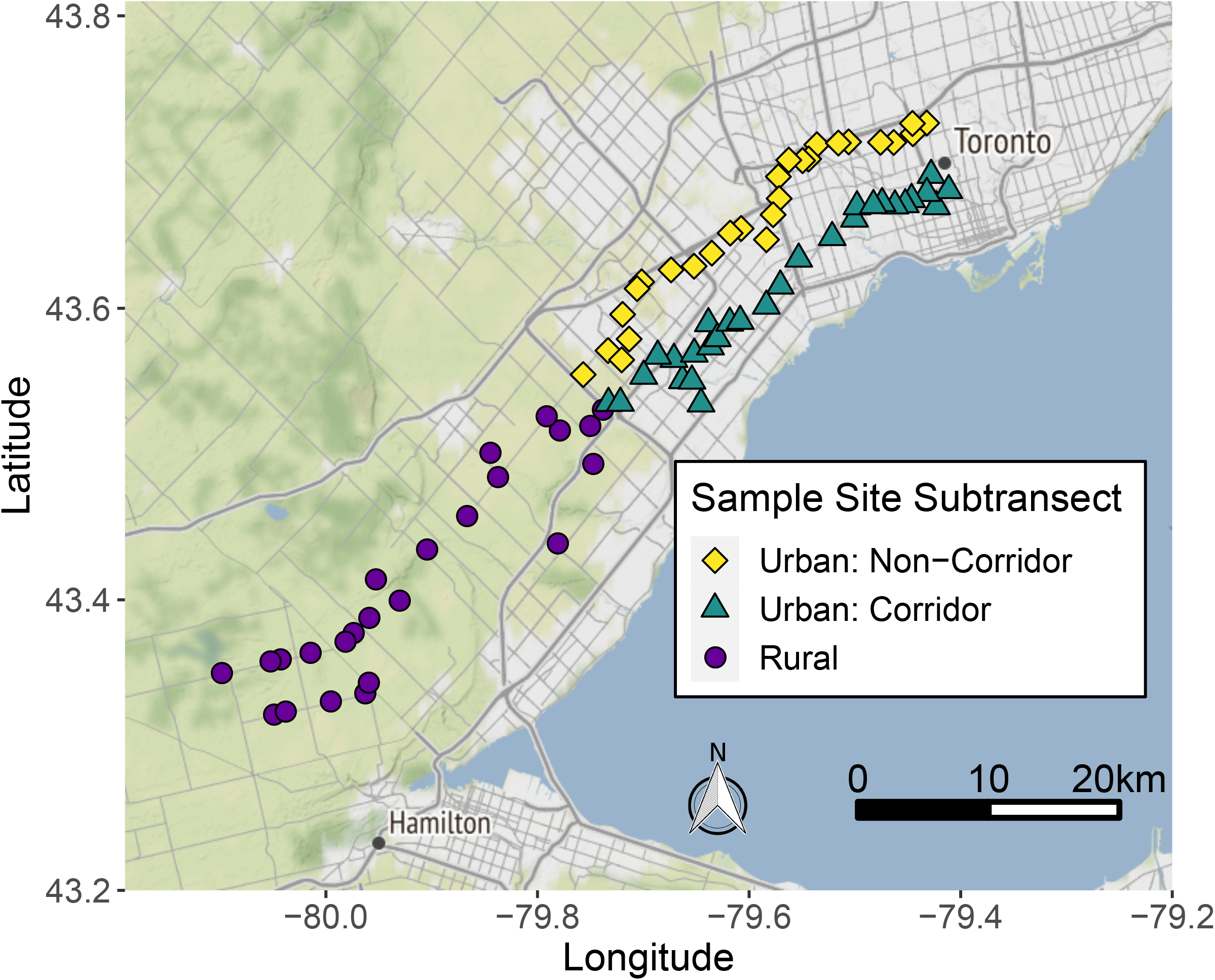
Map of 80 common milkweed populations found along Toronto ‘s urban-rural gradient. Urban: Non-Corridor populations (yellow squares; N= 27). Urban: Corridor populations (green triangles; N=29). Rural populations (purple circles; N=24). The Stamen terrain basemap shows urban and suburban areas in light gray, nonurban agricultural and forested areas in green, and Lake Ontario in blue. Map tiles by Stamen Design, under CC BY 3.0. Data by OpenStreetMap, under ODbL

We surveyed plant reproductive investment and success in 2018 and 2019, plus pollinator visitation in 2019. Each day we visited ca. 10 neighboring populations. To minimize bias due to changes in weather between sampling days, we randomized the order by which sets of ca. 10 populations were visited with regard to subtransect and distance to the urban center. We recorded the latitude and longitude coordinates of each population with a handheld GPS. We then divided the population into equally-sized sections (five for plant reproductive investment/success, three for pollinator visitation) and haphazardly sampled one ramet per section, aiming to separate ramets by >3 m between ramets to avoid resampling the same clone.

We estimated male fitness during peak flowering in July 2018 and 2019. Male fitness per plant was estimated as the mean number of missing pollinaria from nine flowers per ramet, selected from three haphazardly-selected flowers from each of three inflorescences (fewer if less were available). Several studies of *A. syriaca* showed that the number of pollinaria removed is correlated with the number of seeds sired, whereby the number of pollinaria removed sets the upper limit of a plant ‘s male fitness (Broyles & Wyatt 1990; LaRosa 2015).

In July 2019, we also conducted pollinator surveys on three flowering *A. syriaca* ramets per population. We sampled fewer ramets when less than three ramets were flowering in a population. Sampling occurred during periods of high pollinator activity (9:30-17:00) under optimal weather conditions (i.e., <50% cloud cover, no precipitation, temperature >15°C, wind <20 km/h) at populations with flowering *A. syriaca* (58 sites). Pollinators were observed with binoculars from 3 m away for 5 minutes per ramet. The abundance of each pollinator was recorded and species were identified to taxonomic species or morphospecies based on physical descriptions.

In September 2018 and 2019, we measured reproductive effort and female fitness from five ramets from the same populations described above. Reproductive effort in flowering was measured as the number of inflorescences produced during the entire summer. Female fitness was quantified as the total number of non-aborted follicles produced. Fruit set was measured as the mean number of non-aborted follicles per inflorescence. We also measured ramet height as a surrogate of ramet biomass (r = 0.95, N = 31; Online Resource 2).

### Urbanization metrics

To examine the effects of urbanization and green corridors on plant reproduction and pollinators, we quantified urbanization using two methods. First, we determined the distance from each population to the Toronto urban center (43.656327, −79.380904) with the “distHaversine” function from the *geosphere* R package (version 1.5.14, Hijmans 2021). Distance from the urban center along this urban-rural transect is well-correlated with multiple metrics of urbanization such as impervious surface and building density and has been shown to influence ecology and evolution in other plant, herbivore, and pollinator communities (Johnson et al. 2018; Rivkin et al. 2020; Murray-Stoker & Johnson 2021). Hence, we refer to the transect as an urban gradient throughout this paper and use distance from the urban center as a proxy for the degree of urbanization. Second, we used the *UrbanizationScore* software (Czúni et al. 2012; Seress et al. 2014; Lipovits et al. 2015) to calculate a local urbanization score for each population based on the quantity of vegetation, buildings and paved roads within a 1km radius of the site at 100m resolution (Online Resource 1, Figure S-1).

### Statistical analyses

#### Question 1: How does urbanization influence plant reproduction?

To determine how multiple estimates of reproductive success, reproductive effort, fruit set, and plant productivity were affected by urbanization, we fitted general linear mixed-effect models. Models were fitted using maximum likelihood with the *lme4* (version 1.1.21, Bates et al. 2015), *lmerTest* (version 3.1.1; Kuznetsova et al. 2017), and *glmmTMB* (version 1.0.2.1; Brooks et al. 2017) packages. For each response variable (i.e., pollen removal, follicle count, inflorescence count, mean no. of follicles per inflorescence, and height), we used the following full model:

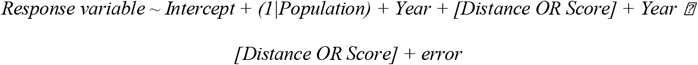

In this model, (1|Population) indicates that population was treated as a random effect. Year was treated as a fixed effect, and distance from the urban center (Distance) and urbanization score (Score) were considered continuous fixed effects. For both Distance and Score models, we fit the following distributions to the data: pollen removal – Poisson distribution; follicle and inflorescence counts subtracted 1, then used zero-inflated, truncated Poisson distributions to account for an excess of ones. We used the “lmer” function from the *lme4* package for the general linear mixed models (mean no. of follicles per inflorescence) and the “glmmTMB” function from the *glmmTMB* package for the remaining generalized linear mixed models because this package can fit zero-inflated models. We used *glmmTMB* for the pollen removal variable because the model would not converge with *lme4* unless “Year” was removed and separate models were run for each of two years (2018, 2019). These two models yielded identical results to the single *glmmTMB* model.

We then fitted reduced models without the interaction term (Year ⍰ [Distance OR Score]), ranked all full and reduced models based on Akaike ‘s information criterion (AIC; Bozdogan 1987), selected the model with the lowest AIC as the best model, and retained statistically equivalent models within 2 ΔAIC of the best model (Online Resource 1, Table S-6). We used the “Anova” function from the *car* package (version 3.0.9; Fox & Weisberg 2019) to perform ANOVA with type III sums-of-squares and the Kenward–Roger method for denominator degrees of freedom approximation to test the significance of the fixed effects (Kenward & Roger 1997; Landsheer & van den Wittenboer 2015). If models without interactions were the best fitting models then we removed the interaction and reran the analyses with type II sums-of-squares (Langsrud 2003).

We inspected model diagnostics with the *DHARMa* package (version 0.4.5, Hartig 2022) and transformed variables to meet the assumptions of normality and homogeneity of variance when necessary (Online Resource 1, Tables S-1–S-4). We explored alternative fitting models based on the change in AIC, and while we show the optimal model, alternative models within 2 ΔAIC typically gave identical conclusions (Online Resource 1, Tables S-6– S-9).

#### Question 2: How do green corridors in urban and suburban areas affect plant reproduction?

To determine how green corridors versus the urban matrix influenced the same response variables among only urban populations (“Urban: Corridor” and “Urban: Non-Corridor”), we fitted general linear mixed-effect models with subtransect added as a fixed effect using the same packages described above. We fitted the following full model:

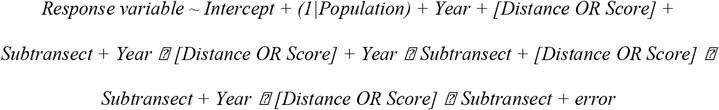

We used identical data distributions for these response variables as detailed in Question 1. We used “lme4” for the mean no. of follicles per inflorescence and pollen removal models, then “glmmTMB” for the remaining models. We then fitted and compared nine nested models using AIC. Briefly, we included the following models in our candidate set: Model 1 = full model; Model 2 = removed third-order interaction term; Models 3-5 = main effects and two second-order interactions; Models 6-8 = main effects and one second-order interaction; and Model 9 = main effects without second-or third-order interactions. We then repeated the post-model-fitting procedures as detailed for Question 1 (Online Resource 1, Table S-7).

#### Question 3: Does the abundance, diversity, and richness of urban pollinators of A. syriaca change along an urbanization gradient?

To assess changes in pollinator communities visiting *A. syriaca* populations along the urbanization gradient, we fitted linear models using the *lme4, lmerTest*, and *car* packages. We regressed each response variable (i.e., pollinator abundance, diversity, and richness) with either distance from the urban center (Distance) or urbanization score (Score) as a fixed effect. Pollinator abundance was calculated as the mean number of pollinators observed per plant, per population. Pollinator diversity was estimated as the inverse of Simpson ‘s diversity index (1/D) using the “diversity” function from the *vegan* package (version 2.5.6; Oksanen et al. 2019). To weight species richness by sampling effort, pollinator richness was calculated as per Menhinick ‘s richness index (Menhinick 1964): the number of species divided by the square root of the number of individuals sampled. We fitted the following full model and repeated the post-model-fitting procedures as detailed for Question 1:

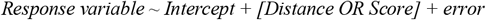

#### Question 4: How do green corridors in urban and suburban areas affect pollinators?

To determine how green corridors versus the urban matrix influenced the same pollinator response variables among only urban populations (“Urban: Corridor” and “Urban: Non-Corridor”), we fitted linear models. For these models, subtransect type was added as a second predictor to the model detailed in Question 3. We fitted the following full model, then fitted a reduced model without the interaction term and repeated the post-model-fitting procedures as detailed for Question 1:

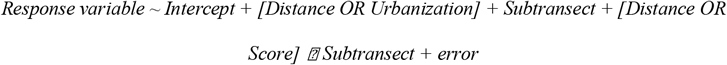

All above analyses were performed using R v.3.5.1. (R Core Team, 2020).

## Results

### Question 1: How does urbanization influence plant reproduction?

Urbanization influenced multiple aspects of plant reproduction. Urbanization reduced male fitness (i.e., pollinaria removed), although the magnitude of this effect varied between years. Within a given year, the number of pollinaria removed from flowers was negatively impacted by urbanization such that there were on average 20% (2018) and 11% (2019) of pollinaria removed per flower at the urban center compared to 39% (2018) and 38% (2019) at the furthest rural populations (Figure 2A). Female fitness (i.e., no. of follicles) was influenced by urbanization differently each year such that the most urban populations produced 11% more follicles in 2018 relative to the furthest rural populations, but 4% fewer in 2019 (Figure 2B). Urbanization positively influenced fruit set (i.e., mean no. of follicles per inflorescence) such that there were 28% (2018) and 29% (2019) more follicles per inflorescence at the urban center versus the furthest rural populations (Figure 2C). Urbanization did not influence reproductive effort (i.e., no. of inflorescences; Figure 2D). Results for an alternative metric of urbanization (urbanization score) were mostly consistent with effects of distance and are presented in the supplement, along with results for plant productivity (i.e., plant height; Online Resource 1, Table S-6).

**Fig. 2.**
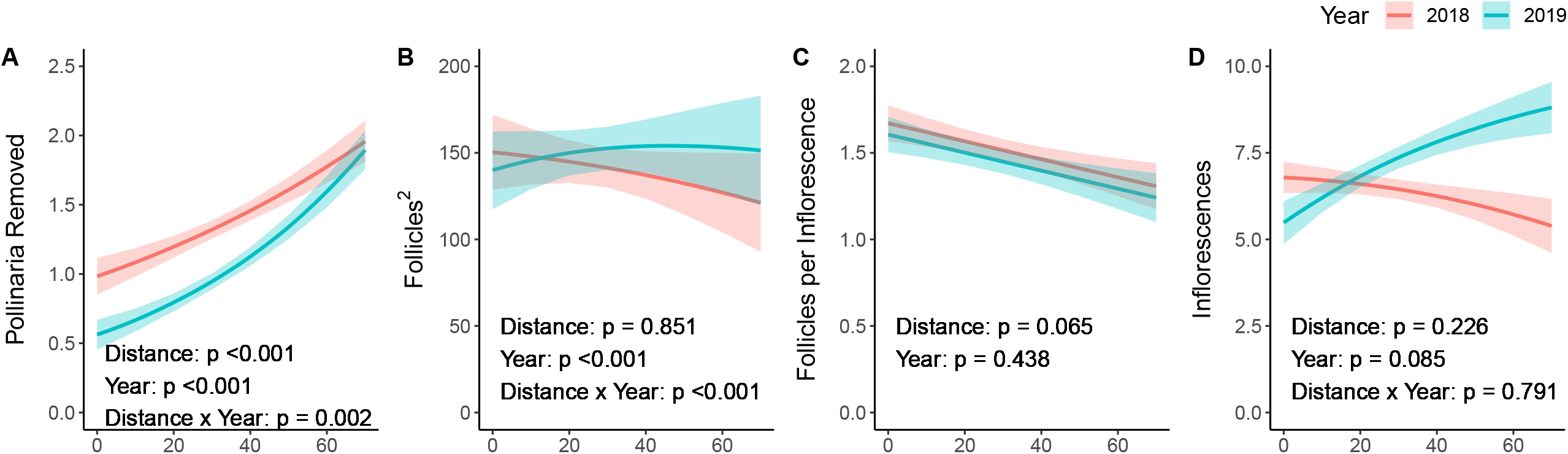
The effects of year and urbanization on pollinaria removed, follicles, mean no. of follicles per inflorescence, and inflorescences for 2018 and 2019 sampling seasons when urbanization was quantified by distance from the urban center. Lines are smoothed curves that represent fitted means (± standard error) are shown for general and generalized linear mixed effects models

### Question 2: How do green corridors in urban and suburban areas affect plant reproduction?

Proximity to the urban corridor influenced male fitness and reproductive effort but not female fitness. The extent to which male fitness (i.e., pollinaria removed) changed with urbanization differed between subtransects, and the magnitude of difference depended on the year (Table 1A). In 2018, urbanization decreased pollen removal such that there was an 83% and 34% decrease in average pollinaria removed per flower from the furthest suburban populations to the urban center along the non-corridor and corridor subtransects respectively, but in 2019, this trend reversed such that there was a 77% and 28% increase in pollen removed with urbanization along the non-corridor and corridor, respectively (Figure 3A). Fruit set (i.e., mean no. of follicles per inflorescence) and female fitness (i.e., no. of follicles) did not vary with subtransect (Table 1B, 2C), although there was an effect of urbanization on fruit set that varied between years as described for the results in Question 1 (Table 1B). Reproductive effort (i.e., no. of inflorescences) was higher along the corridor than the non-corridor subtransect in both 2018 (35%) and 2019 (9%), and decreased with urbanization along the two subtransects such that there were an estimated 34% (2018) and 55% (2019) fewer inflorescences at the urban center compared to the furthest suburban populations (Figure 3D, Table 1D). Results for the alternative metric of urbanization (urbanization score) produced contrasting results for the effects of distance and subtransect and are presented in the supplement, along with results for plant productivity (i.e., plant height; Online Resource 1, Tables S-6 & S-10).

**Table 1.** Results from general and generalized linear mixed models examining the effects of urbanization and proximity to an urban corridor on pollinaria removed, follicles, mean no. of follicles per inflorescence, and inflorescences. Urbanization was quantified via distance from the urban center, and only urban populations were included. Shown are maximum likelihood χ2 and p-values obtained from a type III sums-of-squares ANOVA

|  | Pollinaria Removed |  | Follicles |  | Follicles per Inflorescence |  | Inflorescences |  |
| --- | --- | --- | --- | --- | --- | --- | --- | --- |
| Predictor | $\chi^2$ | $p$ | $\chi^2$ | $p$ | $\chi^2$ | $p$ | $\chi^2$ | $p$ |
| Distance to City Center | 5.706 | 0.017* | 0.974 | 0.324 | 3.349 | 0.067 | 4.556 | 0.033* |
| Year | 6.788 | 0.009** | 172.145 | <0.001*** | 9.128 | 0.003** | 1.966 | 0.161 |
| Subtransect | 5.041 | 0.025* | 0.67 | 0.413 | 0 | 0.987 | 3.82 | 0.051 |
| Distance to City Center x Subtransect | 2.492 | 0.114 | - | - | - | - | 0.031 | 0.86 |
| Distance to City Center x Year | 11.130 | <0.001*** | 0.247 | 0.619 | 9.896 | 0.002** | 0.259 | 0.611 |
| Year x Subtransect | 7.202 | 0.007** | 0.324 | 0.569 | - | - | 2.398 | 0.121 |
| Distance to City Center x Year x Subtransect | 4.622 | 0.032* | - | - | - | - | 0.187 | 0.665 |

**Fig. 3.**
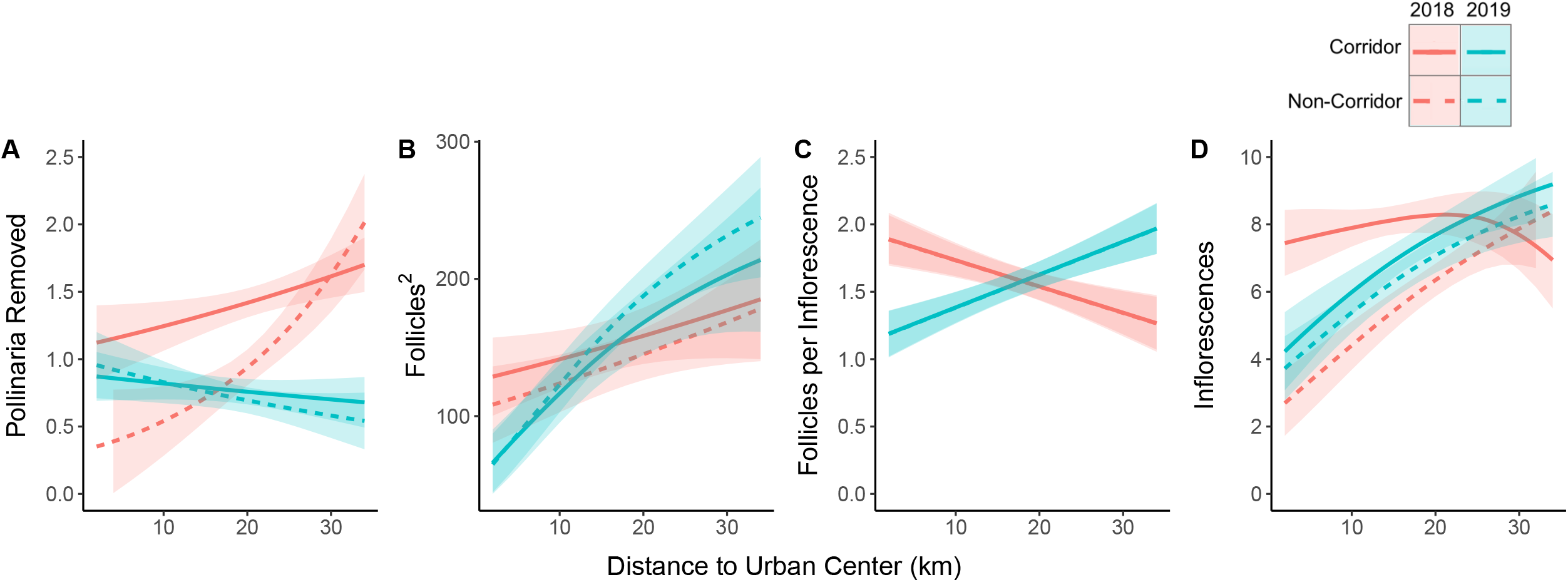
The effects of year, subtransect, and urbanization on pollinaria removed, follicles, mean no. of follicles per inflorescence, and inflorescences for the 2018 and 2019 sampling seasons when urbanization was quantified by distance from the urban center. Lines are smoothed curves representing fitted means (± standard error) are shown for general and generalized linear mixed effects models. Panel C includes 4 overlapping lines

### Question 3: Does the abundance, diversity, and richness of urban pollinators of A. syriaca change along an urbanization gradient?

Urbanization influenced pollinator abundance and species richness but not Simpson ‘s diversity. Abundance was negatively associated with urbanization such that there were an estimated 61% fewer pollinators observed at the urban center versus the furthest rural populations (Figure 4A). Conversely, species richness was 63% higher in the urban center relative to the furthest rural populations (Figure 4B). Urbanization did not influence Simpson ‘s diversity (Figure 4C). Results for the alternative metric of urbanization (urbanization score) were mostly consistent for effects of distance (Online Resource 1, Table S-8).

**Fig. 4.**
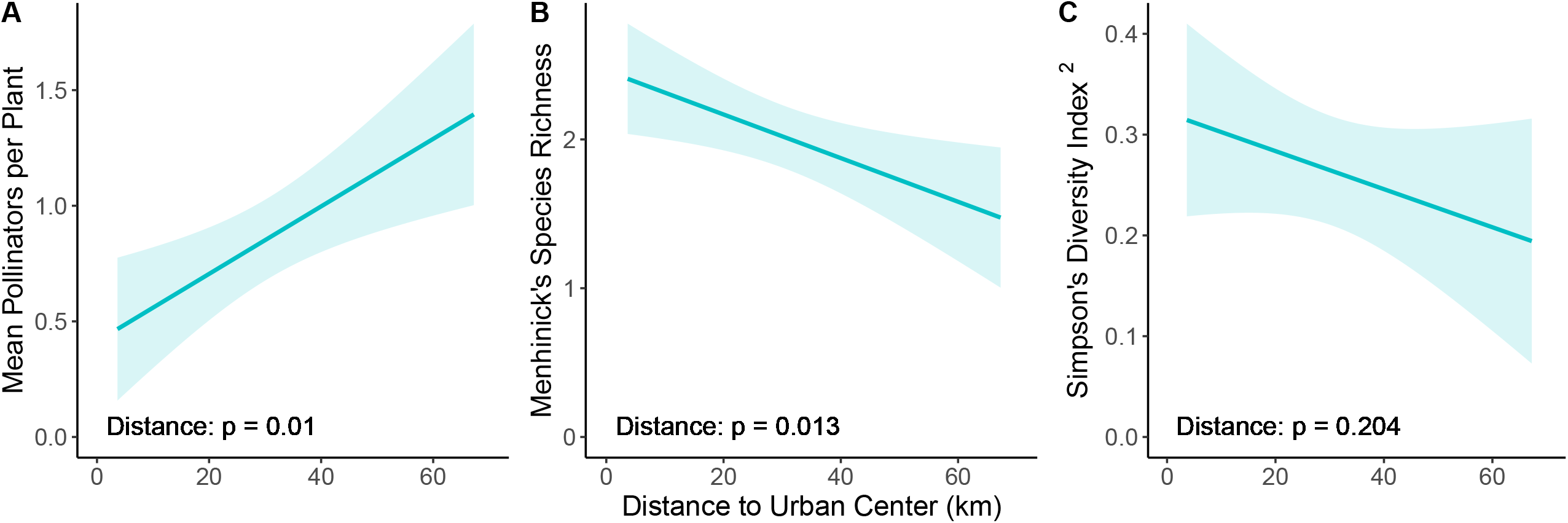
The effects of urbanization on pollinator abundance, Menhinick ‘s species richness, and Simpson ‘s diversity for the 2019 sampling season when urbanization was quantified by distance from the urban center. Lines of best fit (± standard error) are shown for linear models

### Question 4: How do green corridors in urban and suburban areas affect the abundance, diversity, and richness of urban pollinators of A. syriaca?

Green corridors did not influence pollinator abundance but did affect species richness and Simpson ‘s diversity. While green corridors and urbanization did not affect pollinator abundance (Figure 5A), species richness and Simpson ‘s diversity were both negatively associated with green corridors such that they were 21% and 76% lower along the corridor compared to the non-corridor subtransect, respectively (Figure 5B-C). Results for the alternative metric of urbanization (urbanization score) were mostly consistent for effects of distance (Online Resource 1, Table S-9).

**Fig. 5.**
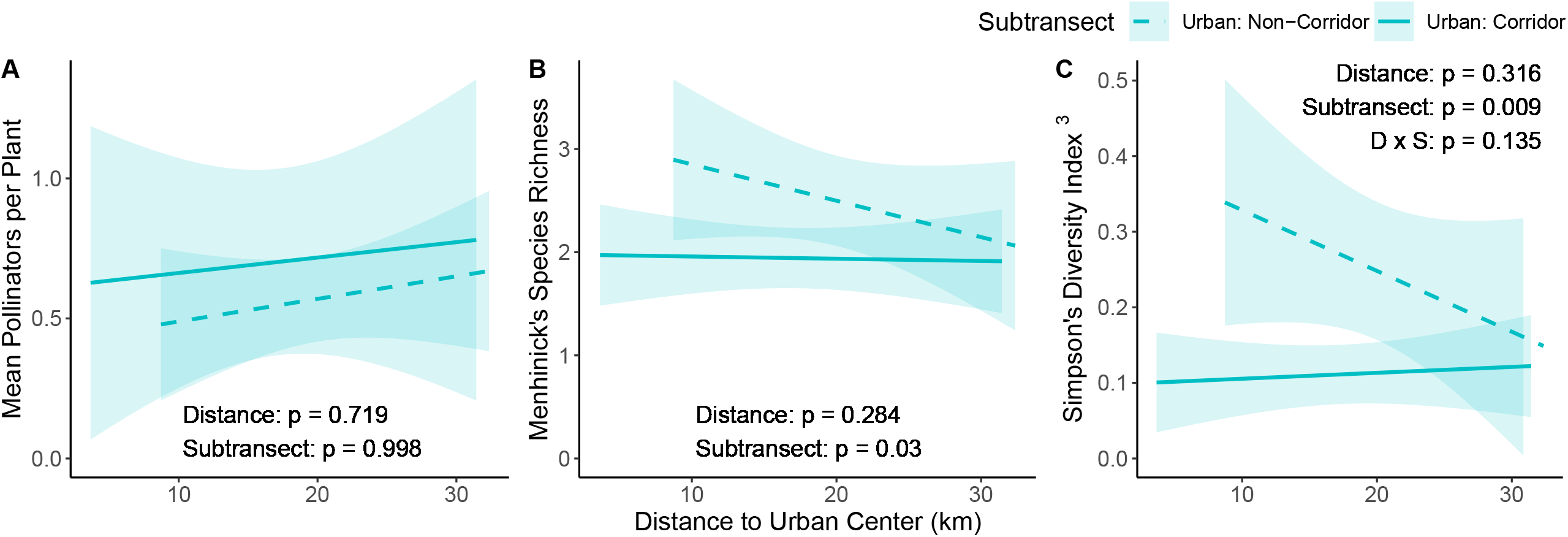
The effects of subtransect and urbanization on pollinator abundance, Menhinick ‘s species richness, and Simpson ‘s diversity for the 2019 sampling season when urbanization was quantified by distance from the urban center. Lines of best fit (± standard error) are shown for linear models

## Discussion

In this study, we tested the hypothesis that urban environments and green corridors influence plant reproduction by studying natural populations of an obligately outcrossing plant along an urbanization gradient. Our results show that both urban environments and green corridors influence reproduction in common milkweed. Four key results are essential for answering our research questions. First, urbanization influenced multiple aspects of reproduction in common milkweed: male fitness (i.e, pollen removal), female fitness (i.e., no. of follicles), and fruit set (i.e., mean no. of follicles per inflorescence). Second, proximity to a green corridor influenced only male fitness and reproductive effort (i.e., no. of inflorescences). Third, urbanization affected pollinator abundance and species richness but not Simpson ‘s diversity. Fourth, proximity to a green corridor influenced pollinator species richness and Simpson ‘s diversity but not abundance. Overall, these results demonstrate that urbanization, green corridors, and pollinator communities can shape the reproductive investment and fitness of natural plant populations in urban environments.

### How does urbanization influence plant reproduction and urban pollinators?

There is no consensus about how urbanization impacts plant reproduction (Cheptou & Avendaño 2006; Van Rossum 2010; Pellissier et al. 2012; Verboven et al. 2012) or pollinators (Banaszak-Cibicka & Żmihorski 2012; Theodorou et al. 2016; Wenzel et al. 2019; Pereira et al. 2021; Prendergast et al. 2022). Similarly, urbanization has been found to influence plant-pollinator interactions in highly variable and contrasting ways (Harrison & Winfree 2015). Despite considerable variation in the specific urban drivers of altered plant-pollinator interactions, plants and pollinators are expected to experience parallel responses to urbanization due to their mutualistic interdependence within plant-pollinator networks (Harrison & Winfree 2015; Rivkin et al. 2020). Our results partially support this prediction. Urbanization had varying effects on reproductive success of *A. syriaca* depending on the component of plant reproduction studied. For example, urbanization consistently decreased male fitness (pollen removal) yet influenced female fitness (no. of follicles) negatively in 2018 and positively in 2019. In urban areas, pollinator abundance declined while species richness increased and Simpson ‘s diversity did not change. Since Menhinick ‘s species richness accounts for the number of individuals observed, species richness was not reduced proportionally with abundance. The complexity of these results are likely due to several biotic and abiotic aspects of urban environments that vary over space and time.

Our observation that urbanization did not influence plant reproductive effort contrasts with other studies of urban plant populations. Existing literature suggests urbanization lowers plant reproductive effort: Experimental populations of *Brassica rapa* produced fewer flowers, and natural populations of *Paubrasilia echinata* produced fewer flowers per inflorescence, in urban vs. rural environments (Oliveira et al. 2019; Rivkin et al. 2020). Our results suggest that urbanization does not influence plant reproductive success in *A. syriaca* and that plants experience relatively consistent growth and development— a view corroborated by consistent plant productivity (height) along the urban-rural gradient. Plants in the populations closest to the urban center experienced lower male fitness (pollen removal), however, and were visited by fewer individual pollinators compared to the most distant rural populations. This result aligns with previous work that identified altered plant-pollinator interactions as a major contributor to depressed reproductive success in urban plants (Cheptou & Avendano 2006; Pellissier et al. 2012; Geslin et al. 2013). Conversely, our finding that the effects of urbanization on reproductive success can vary over time has been shown, to our knowledge, in only one other study (Rivkin et al. 2020). A 19% decrease in total precipitation during the 2019 growing season (April-September) compared to 2018 (Online Resource 1, Figure S-6) may have induced physiological stress that could explain why urban plants displayed higher female fitness in 2018 but lower in 2019. Lastly, the relatively high fruit set experienced by urban plants has been observed in plants capable of self-fertilization (Ushimaru et al. 2014). However, given that *Asclepias syriaca* is largely self-incompatible (Wyatt 1994), it is possible that this result reflects increased non-self pollen deposition per inflorescence due to more pollinator visits per plant in small urban populations. The higher pollinator species richness in urban areas may also reflect more diversity in pollination behavior; despite lower abundance, urban pollinators may have alleviated pollen limitation by delivering mostly outcrossed pollen and thus increasing fertilization rates in *A. syriaca*. In contrast, pollinators like *Apis mellifera* and *Bombus spp*. can effect relatively high rates of geitonogamy in *Asclepias spp*. (Ivey et al. 2003; Howard & Barrows 2014). Future studies should assess whether urbanization impacts self-pollination rates in *A. syriaca*.

Our results demonstrate that urban environments have complex influences on plants, pollinators, and the ecological interactions between them. This work diverges from previous studies due to its focus on how multiple components of plant-pollinator interactions are shaped by urbanization: i.e., the separate ecologies of the plants and pollinators as well as the consequences of the species interactions for plant fitness. Our results suggest that while urbanization may impact plant-pollinator interactions, the long-term fitness consequences for the host plants are not straightforward. Specifically, even though we found that aspects of both male and female fitness were affected by urbanization, we did not integrate sexual and vegetative reproduction over the entire lifetime of a plant and thus quantify total fitness. Future studies could advance our understanding of the effects of urbanization on plants by measuring plant fitness over longer periods of time. It would also be prudent to test for differences in pollen and seed quality, measured by pollen grain count (Wyatt et al. 2000) or possibly size distribution (Kelly et al. 2002), as well as seed germination success and seedling survival (Wen & Simons 2020).

### How do green corridors in urban and suburban areas affect plant reproduction and urban pollinators?

Little is known about how corridors influence plant-pollinator interactions in urban environments. In non-urban areas, however, corridors in fragmented habitats have often been found to facilitate pollination through increased pollinator movement and pollen transfer (Tewksbury et al. 2002; Townsend & Levey 2005). As a result, plants growing within or near corridors frequently experience increased pollination and/or reproductive success (Cranmer et al. 2012; Kormann et al. 2016). Our study represents one of the first examinations of how corridors affect plant reproduction in urban environments. We found that proximity to an urban corridor influenced male fitness differently each year, consistently increased reproductive effort, but decreased pollinator species richness and Simpson ‘s diversity. These complex results indicate that the mechanisms which typically benefit plant-pollinator interactions in corridors may be rearranged by the various interacting aspects of urban environments.

In 2018, urban plants growing within or near the corridor experienced significantly higher male fitness and reproductive effort than urban plants growing far from the corridor. These divergent responses may signal that corridors buffered plant-pollinator interactions from negative consequences of urban development such as impervious surfaces and soil compaction, which can induce plant stress by reducing water and nutrient absorption (Oke et al. 1989; Lipiec & Stępniewski 1995). The green corridor did not significantly influence female fitness or fruit set, suggesting that green corridors may exert the most impact on plant reproductive processes that occur in the earlier portion of the growing season when inflorescences are developing or when flowers open and are pollinated. While our ability to contextualize these observations with our pollinator data is limited due to pollinator sampling in 2019 only, we found that plant reproductive success and effort was relatively consistent among corridor and non-corridor subtransects in 2019, whereas pollinator species richness and Simpson ‘s diversity were lower along the corridor subtransect. Assuming similar milkweed abundance among the subtransects, these results could signal that the corridor subtransect ‘s high edge density (ratio of patch edge to area; McGarigal & Marks 1995) dampened pollinator diversity via edge effects (Chisholm et al. 2011) or that lower plant diversity in the corridor reduced the subtransect ‘s pollinator diversity (Theodorou et al. 2020b). However, we speculate that milkweed populations within the corridor subtransect contained more plants than the non-corridor subtransect populations and consequently attracted fewer pollinators on a per-plant level. This scenario would represent a dilution effect wherein a similar or even higher quantity of pollinators in corridors would be distributed among more corridor-dwelling plants and give the appearance of lower richness and diversity per plant. A general plant-pollinator dilution effect probably co-occurred wherein the corridor subtransect contained more plants that share the same main pollinators of common milkweed, i.e. generalists such as *A. mellifera* and *Bombus spp*. (Kephart 1983; MacIvor et al. 2017).

Our study is among the first to demonstrate that urban corridors can impact pollinator community structure and plant reproduction. While plant-pollinator interactions may have differed in the green corridor, we did not observe consistent, marked differences in plant reproductive success among the subtransects. These results are consistent with findings that species respond variably to corridors (Resasco 2019), and suggest that linear green corridors may play a relatively minor, or at least cryptic, role in facilitating plant-pollinator interactions in urban environments in this system. Future studies should assess how green corridors influence the plant offspring quality by investigating the genetic consequences of urban corridors on plant reproduction. More broadly, it is essential to understand how urban corridors influence interactions among species in general.

## Conclusions

Within the past few decades, ecologists have made great strides in resolving how urban environments influence the ecology of populations and communities. However, it remains unclear how urbanization affects the interactions among species, as well as their consequences for individuals and populations (Shochat et al. 2006; Turrini et al. 2016). Moreover, we do not understand how elements of the heterogeneous urban landscape, such as habitat corridors, affect these interactions. Here, we show that both urban environments and the landscape features within them, specifically green corridors, can restructure plant-pollinator interactions and impact plant reproduction in diverse and sometimes unexpected ways. These results underscore how the spatiotemporal heterogeneity of urban environments can influence species interactions that yield unpredictable consequences for the species involved, a pattern that echoes the vast research attesting to the complexity with which pollinators respond to urbanization (Harrison & Winfree 2015; Wenzel et al. 2019; Prendergast et al. 2022). Green corridors may function differently for pollinators in urban areas; as such, urban planners and conservation groups should consider various tools, in addition to green corridors, for promoting the biodiversity and stability of urban plant-pollinator communities. To maintain ecosystem functioning in the wake of unprecedented levels of global anthropogenic change, further research should utilize integrative methodologies that seek to capture how the biotic and abiotic elements of urban environments shape species interactions and their consequences.

## Acknowledgements

We thank L. Miles for support coordinating field work and data collection; S. Munim and V. Nhan for assistance with field work; J.S. MacIvor for guidance analyzing the pollinator ecology data; the EvoEco and Wagner labs, especially L. Albano, A. Filazzola, D. Murray-Stoker, J. Santangelo, and F. Torres Vanegas, whose comments have greatly strengthened this manuscript. We also wish to acknowledge this land on which the University of Toronto operates. For thousands of years it has been the traditional land of the Huron-Wendat, the Seneca, and the Mississaugas of the Credit. Today, this meeting place is still the home to many Indigenous people from across Turtle Island and we are grateful to have the opportunity to work on this land. It is a privilege for us to perform research on a plant which has traditionally been a source of food, medicine, and fiber for many Indigenous communities, including those who have been caretakers of this land for time immemorial.

## Statements and Declarations

### Funding

This work was funded by an NSERC CREATE Enviro Trainee Grant (SB), NSERC Discovery Grants (HW & MJ), a Canada Research Chair (MJ), and an E.W.R. Steacie Fellowship (MJ).

### Competing Interests

The authors declare no competing interests.

### Author Contribution Statement

SB, MJ, and HW conceived the project. All authors designed the study. SB and AT conducted the field work. SB performed the analyses, wrote the manuscript, and revised it with input from MJ and HW.

### Statement of Data Availability

All data will be submitted to Zenodo and given a DOI upon acceptance of the manuscript.

## Notes

### Competing Interest Statement

The authors have declared no competing interest.

