## Supplemental File 1 for "Urbanization and green corridors influence reproductive success and pollinators of common milkweed"

**Online supplementary for**: Urbanization and green corridors influence reproductive success and pollinators of common milkweed

**Authors**: Sophie Breitbart (1,2,3), Albert Tomchyshyn (2), Helene Wagner (1,2,3), Marc Johnson (1,2,3)

(1) Department of Ecology and Evolutionary Biology

University of Toronto

25 Willcocks Street

Toronto, Ontario

Canada M5S 3B2

(2) Department of Biology

University of Toronto Mississauga

3359 Mississauga Road

Mississauga, ON

Canada L5L 1C6

(3) Centre for Urban Environments

University of Toronto Mississauga

3359 Mississauga Road

Mississauga, ON

Canada L5L 1C6

Contents:

- Supplementary text:
  - Meteorological analysis
- Supplementary tables and figures:
  - Supplementary tables: Tables S1-S10
  - Supplementary figures: Figures S1-S6

**Meteorological analysis**

We used historical weather data to contextualize how yearly differences in precipitation and temperature influenced plant investment and success in reproduction. All data were retrieved from the Digital Archive of Canadian Climatological Data (<https://climate.weather.gc.ca/>). We downloaded data for daily mean temperature and mean precipitation for 2018-2019 from the Oakville weather station (climate ID: 6155750) because it was ca. 5 km from the transect midpoint. We excluded two dates because data were unavailable for both variables.

We then calculated monthly means for temperature and precipitation during 2018-2019 (Figure S-6).

### **Figures**

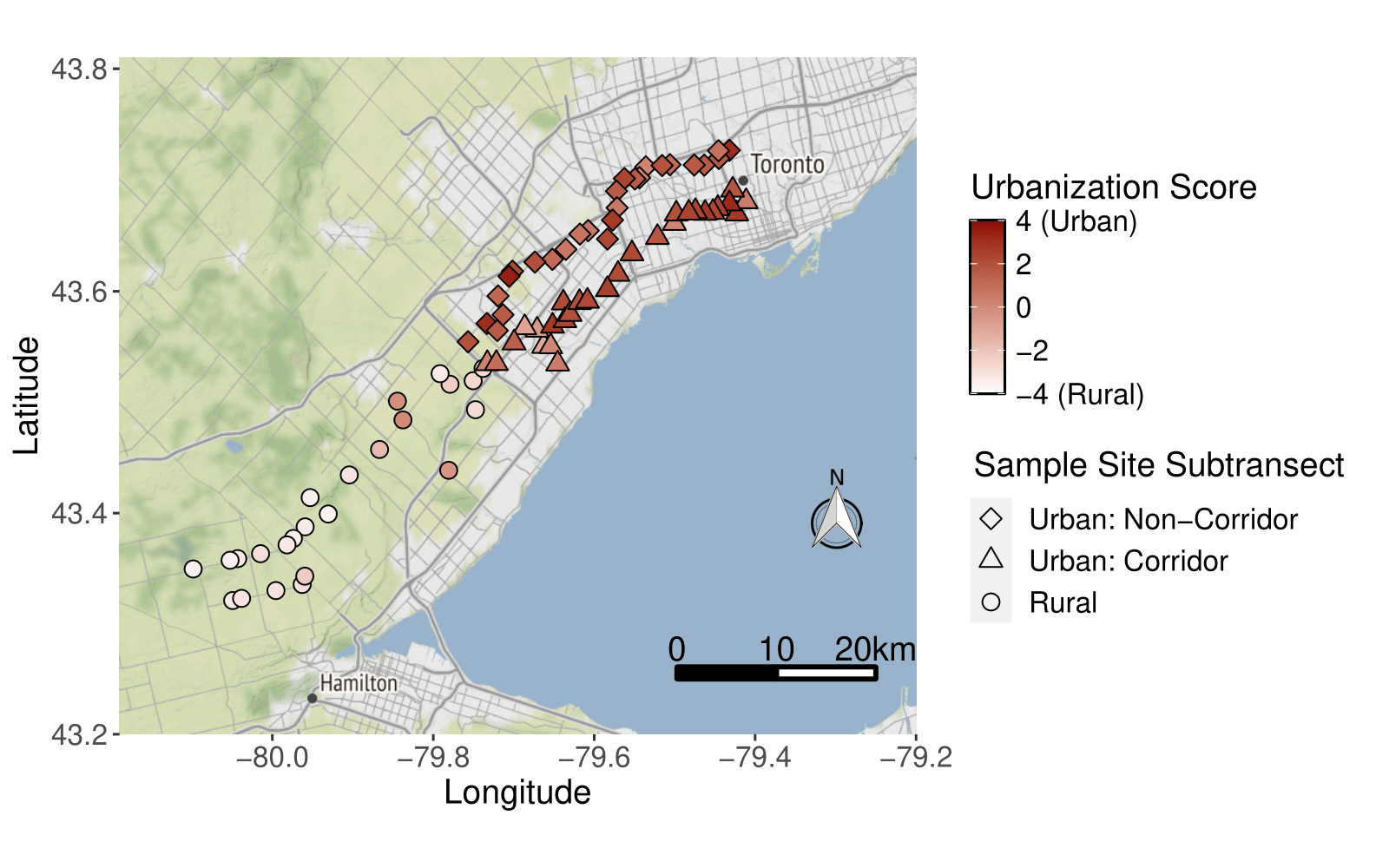

**Fig. S-1** Map of 80 common milkweed populations found along Toronto’s urban-rural gradient. The color of the symbols indicates urbanization score, where positive values indicate a high degree of urbanisation (based on the quantity of vegetation, buildings and paved roads per 1 km^2^). The Stamen terrain basemap shows urban and suburban areas in light gray, nonurban agricultural and forested areas in green, and Lake Ontario in blue. Map tiles by Stamen Design, under CC BY 3.0. Data by OpenStreetMap, under ODbL

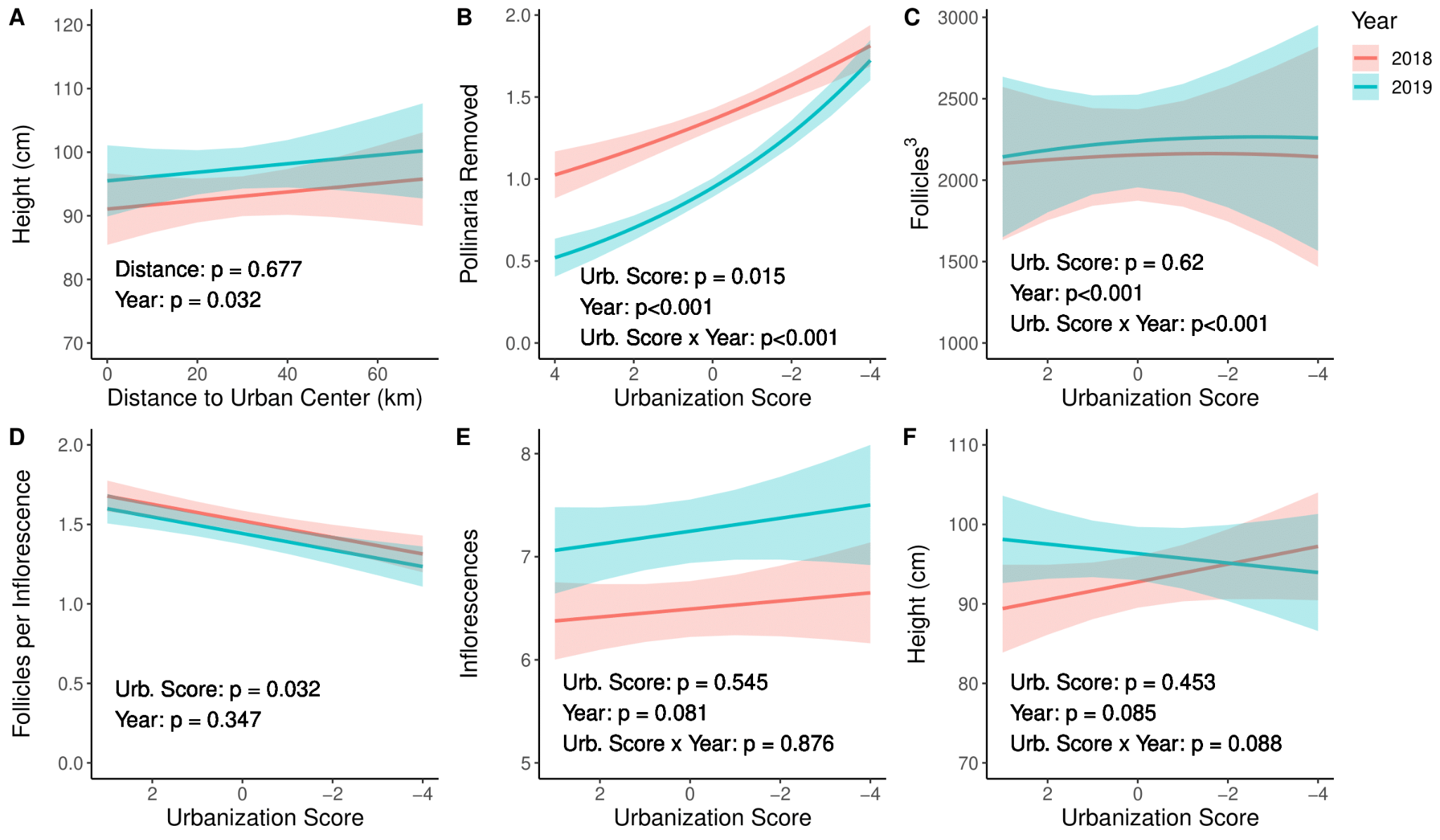

**Fig. S-2** The effects of year and urbanization on pollinaria removed, number of follicles, mean number of follicles per inflorescence, number of inflorescences, and plant height for 2018 (red) and 2019 (blue) when urbanization was quantified by distance from the urban center (A) and urbanization score (B-F). Urbanization score (Urb. Score) spans from 4 (highly urban) to -4 (highly rural). Lines are smoothed curves (± standard error) from general and generalized linear mixed effects models

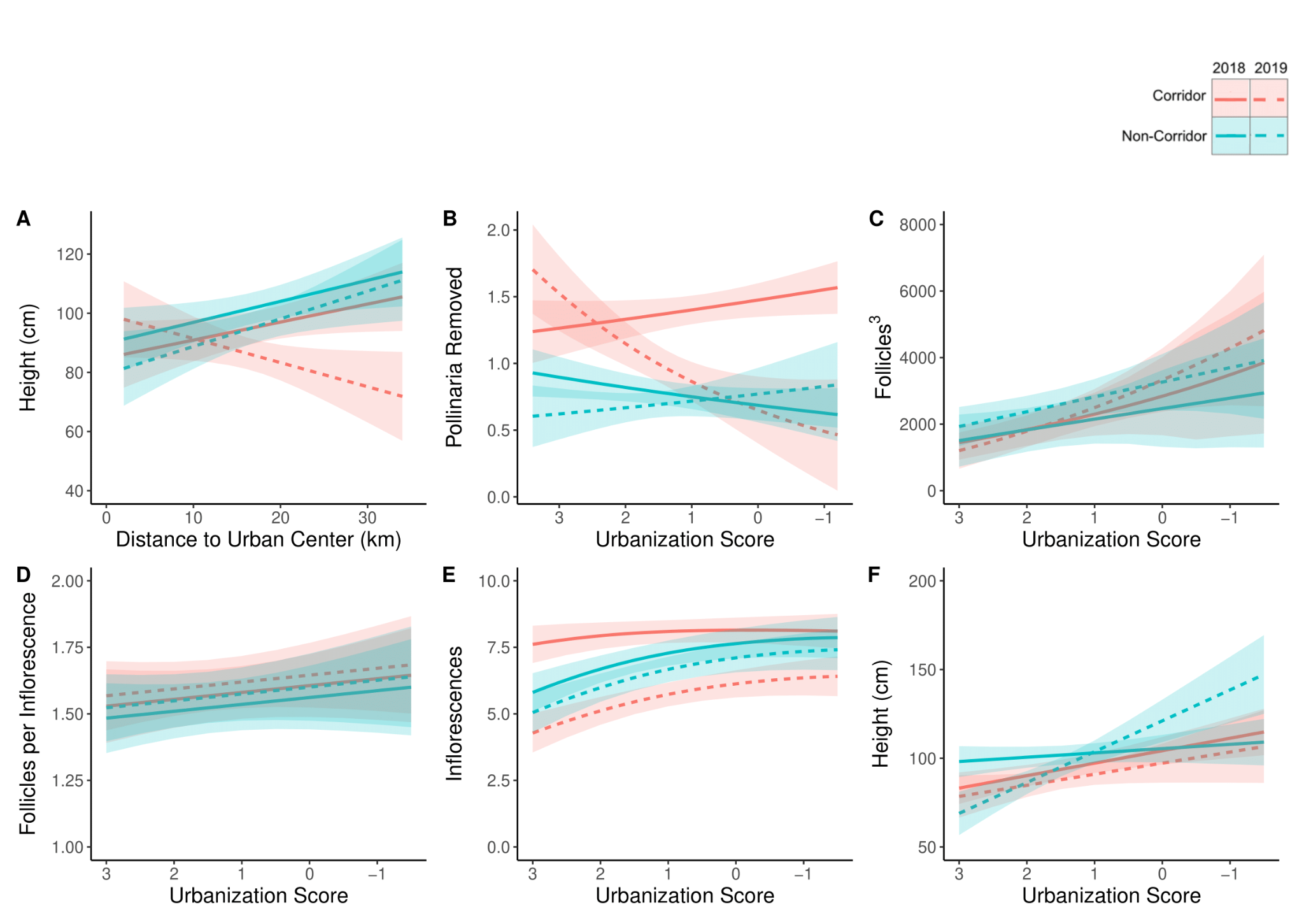

**Fig. S-3** The effects of year, subtransect, and urbanization on plant height (A, F), pollinaria removed (B), number of follicles (C), mean number of follicles per inflorescence (D), and number of inflorescences (E) for 2018 (red) and 2019 (blue) when urbanization was quantified by distance from the urban center (A) or urbanization score (B-F). Urbanization score spans from 4 (highly urban) to -4 (highly rural). Lines are smoothed curves (± standard error) from general and generalized linear mixed effects models. Results from type III sums-of-squares ANOVA are included in Table S-10

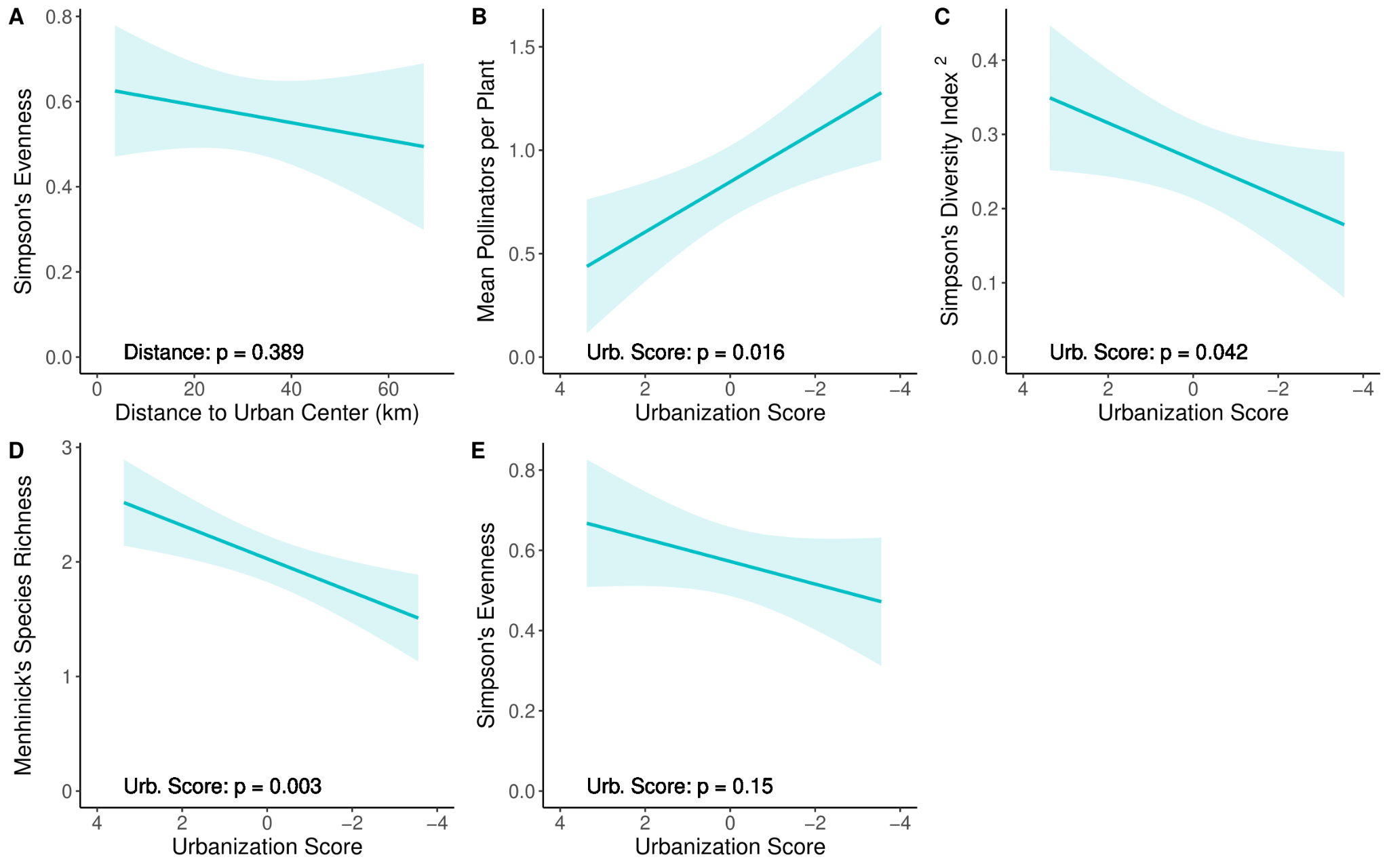

**Fig. S-4** The effects of urbanization on pollinator evenness (A), abundance (B), Simpson’s diversity (C), Menhinick’s species richness (D), and evenness (E) for 2019 when urbanization was quantified by distance from the urban center (A) and urbanization score (B-E). Urbanization score spans from 4 (highly urban) to -4 (highly rural). Lines of best fit (± standard error) are from linear models

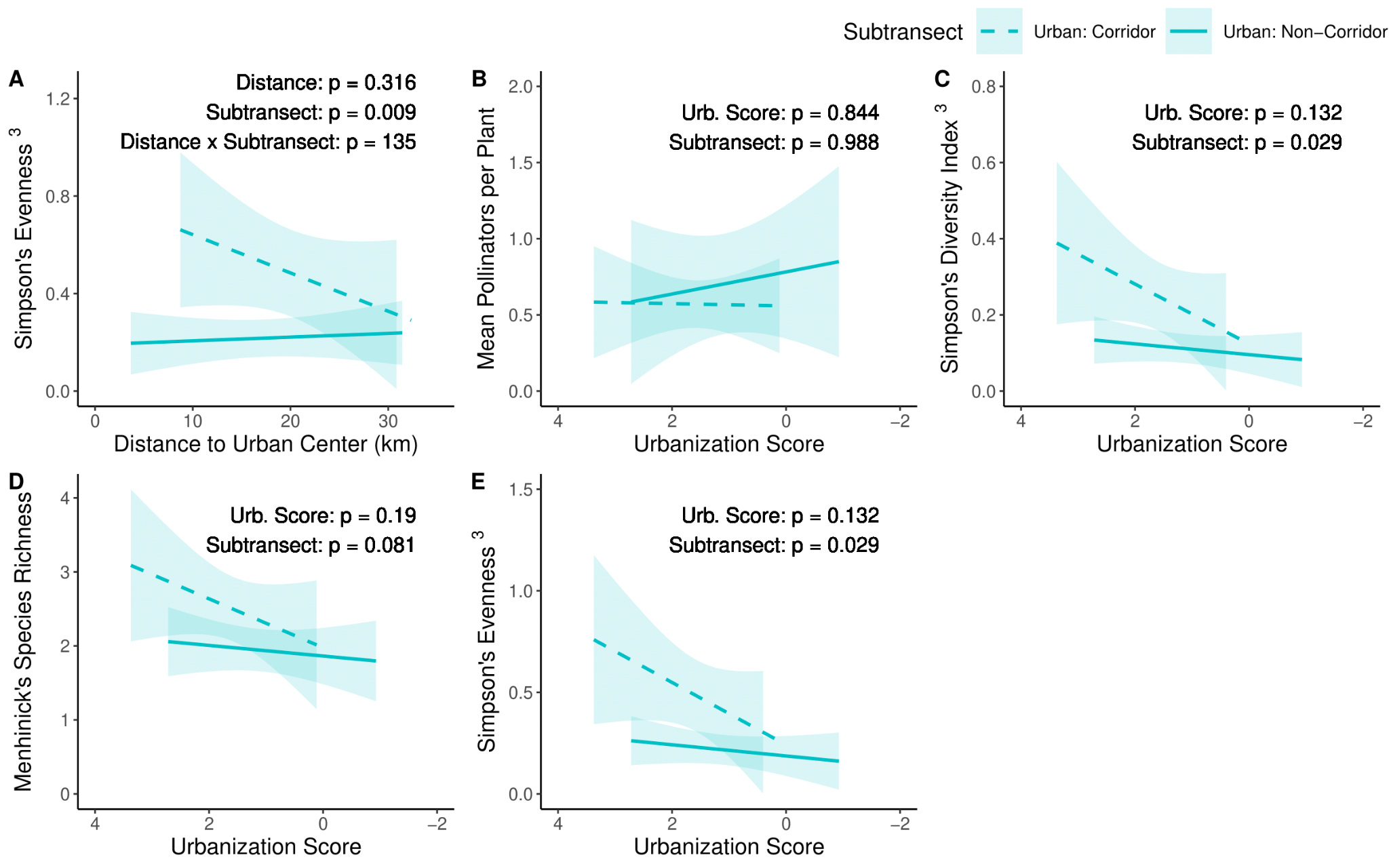

**Fig. S-5** The effects of urbanization and proximity to an urban corridor on pollinator evenness (A), abundance (B), Simpson’s diversity (C), Menhinick’s species richness (D), and evenness (E) for 2019 when urbanization was quantified by distance from the urban center (A) and urbanization score (B-E). Urbanization score spans from 4 (highly urban) to -4 (highly rural). Lines of best fit (± standard error) are from linear models

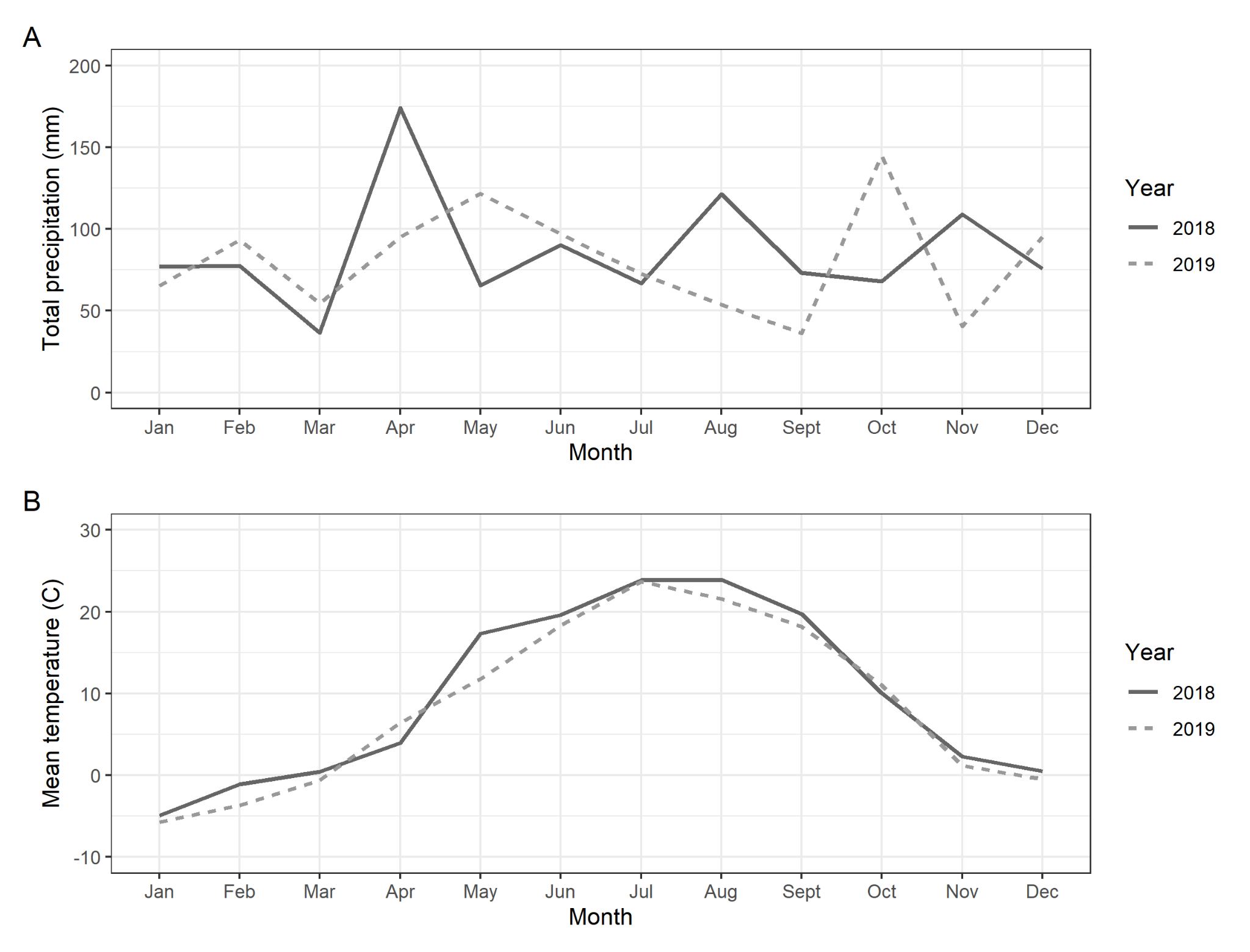

**Fig. S-6** Monthly total precipitation (A) and mean temperatures (B) for 2018-2019 from the Oakville weather station (climate ID: 6155750). Monthly total precipitation and mean temperature were 19% and 8% lower during the 2019 growing season (April-September) compared to the 2018 growing season

### **Tables**

|  | "Distance" Models | | "Urbanization Score" Models | |
| --- | --- | --- | --- | --- |
|  | LMER | GLMER | LMER | GLMER |
| Pollinaria removed | - | None | - | None |
| Follicles | - | x^2^ | - | x^3^ |
| Follicles per Inflorescence | None | - | None | - |
| Inflorescences | - | None | - | None |
| Height | None | - | None | - |

**Table S-1** Data transformations used on plant reproductive success data from all populations to improve normality and homogeneity of variance (Question 1). “X” represents the transformed response variable

|  | "Distance" Models | | "Urbanization Score" Models | |
| --- | --- | --- | --- | --- |
|  | LMER | GLMER | LMER | GLMER |
| Pollinaria removed | - | None | - | None |
| Follicles | - | x^2^ | - | x^3^ |
| Follicles per Inflorescence | None | - | None | - |
| Inflorescences | - | None | - | None |
| Height | None | - | None | - |

**Table S-2** Data transformations used on plant reproductive success data from only urban populations to improve normality and homogeneity of variance (Question 2). “X” represents the transformed response variable

|  | "Distance" Models | "Urbanization Score" Models |
| --- | --- | --- |
| Abundance | None | None |
| Simpson's Diversity | x^2^ | x^2^ |
| Menhinick’s Species Richness | None | None |
| Simpson's Evenness | None | None |

**Table S-3** Data transformations used on pollinator community data from all populations to improve normality and homogeneity of variance (Question 3). “X” represents the transformed response variable

|  | "Distance" Models | "Urbanization Score" Models |
| --- | --- | --- |
| Abundance | None | None |
| Simpson's Diversity | x^3^ | x^3^ |
| Menhinick’s Species Richness | None | None |
| Simpson's Evenness | x^3^ | x^3^ |

**Table S-4** Data transformations used on pollinator community data from only urban populations to improve normality and homogeneity of variance (Question 4). “X” represents the transformed response variable

|  | "Distance" Models | "Urbanization Score" Models |
| --- | --- | --- |
| Pollinaria removed | Poisson | Poisson |
| Follicles | Subtracted 1, then used zero-inflated, "truncated Poisson" family | Subtracted 1, then used zero-inflated, "truncated Poisson" family |
| Inflorescences | Subtracted 1, then used zero-inflated, "truncated Poisson" family | Subtracted 1, then used zero-inflated, "truncated Poisson" family |

**Table S-5** Non-Gaussian probability distributions used on plant reproductive success data from all populations (Questions 1-2)

| **Model** | **Urbanization Metric** | **Best or Alternative Model** | **Variable** | **AIC** | **ΔAIC from Best Model** |
| --- | --- | --- | --- | --- | --- |
| 1 | Distance to Urban Center | Best Model | Pollinaria Removed | 8665.667 | - |
| 2 | Urbanization Score | Best Model | Pollinaria Removed | 8671.383 | - |
| 3 | Distance to Urban Center | Best Model | Follicles | 40409.962 | - |
| 4 | Urbanization Score | Best Model | Follicles | 800665.975 | - |
| 5 | Distance to Urban Center | Best Model | Follicles per Inflorescence | 1404.405 | - |
| 6 | Distance to Urban Center | Alternative Model | Follicles per Inflorescence | 1405.294 | 0.889 |
| 7 | Urbanization Score | Best Model | Follicles per Inflorescence | 1408.854 | - |
| 8 | Urbanization Score | Alternative Model | Follicles per Inflorescence | 1410.854 | 2 |
| 9 | Distance to Urban Center | Best Model | Inflorescences | 2572.89 | - |
| 10 | Urbanization Score | Best Model | Inflorescences | 2574.018 | - |
| 11 | Distance to Urban Center | Best Model | Height | 4713.787 | - |
| 12 | Distance to Urban Center | Alternative Model | Height | 4714.723 | 0.936 |
| 13 | Urbanization Score | Best Model | Height | 4740.032 | - |
| 14 | Urbanization Score | Alternative Model | Height | 4740.879 | 0.847 |

**Table S-6** Summary of AIC scores from general and generalized linear mixed models examining the effects of urbanization on plant reproductive success (Question 1). All alternative models yielded identical conclusions to the best models, although model 14 only retained main effects while the best model retained a marginally significant interaction effect of Urbanization Score x Year (p = 0.088)

| **Model** | **Urbanization Metric** | **Best or Alternative Model** | **Variable** | **AIC** | **ΔAIC from Best Model** |
| --- | --- | --- | --- | --- | --- |
| 1 | Distance to Urban Center | Best Model | Pollinaria Removed | 5028.809 | - |
| 2 | Distance to Urban Center | Alternative Model | Pollinaria Removed | 5029.589 | 0.78 |
| 3 | Urbanization Score | Best Model | Pollinaria Removed | 5043.318 | - |
| 4 | Distance to Urban Center | Best Model | Follicles | 29874.141 | - |
| 5 | Distance to Urban Center | Alternative Model | Follicles | 29874.658 | 0.517 |
| 6 | Distance to Urban Center | Alternative Model | Follicles | 29875.435 | 1.294 |
| 7 | Urbanization Score | Best Model | Follicles | 612252.686 | - |
| 8 | Urbanization Score | Alternative Model | Follicles | 612254.565 | 1.879 |
| 9 | Distance to Urban Center | Best Model | Follicles per Inflorescence | 1037.136 | - |
| 10 | Distance to Urban Center | Alternative Model | Follicles per Inflorescence | 1038.621 | 1.485 |
| 11 | Distance to Urban Center | Alternative Model | Follicles per Inflorescence | 1037.689 | 0.553 |
| 12 | Urbanization Score | Best Model | Follicles per Inflorescence | 1051.574 | - |
| 13 | Urbanization Score | Alternative Model | Follicles per Inflorescence | 1052.593 | 1.019 |
| 14 | Urbanization Score | Alternative Model | Follicles per Inflorescence | 1053.507 | 1.933 |
| 15 | Urbanization Score | Alternative Model | Follicles per Inflorescence | 1053.509 | 1.935 |
| 16 | Distance to Urban Center | Best Model | Inflorescences | 1824.387 | - |
| 17 | Distance to Urban Center | Alternative Model | Inflorescences | 1825.842 | 1.455 |
| 18 | Distance to Urban Center | Alternative Model | Inflorescences | 1825.678 | 1.291 |
| 19 | Urbanization Score | Best Model | Inflorescences | 1838.523 | - |
| 20 | Urbanization Score | Alternative Model | Inflorescences | 1839.223 | 0.7 |
| 21 | Urbanization Score | Alternative Model | Inflorescences | 1839.406 | 0.883 |
| 22 | Distance to Urban Center | Best Model | Height | 3320.556 | - |
| 23 | Distance to Urban Center | Alternative Model | Height | 3321.239 | 0.683 |
| 24 | Urbanization Score | Best Model | Height | 3347.848 | - |
| 25 | Urbanization Score | Alternative Model | Height | 3349.151 | 1.303 |

**Table S-7** Summary of AIC scores from general and generalized linear mixed models examining the effects of urbanization and proximity to an urban corridor on plant reproductive success (Question 2). Alternative models 5, 6, 10, 13-15, and 17 yielded identical conclusions to the best models. Deviations between remaining models were due to marginally significant effects in the best model becoming significant (Model 18) or insignificant (Models 11 & 21) in the alternative model, a significant effect in the best model becoming insignificant (Model 20), or the preservation of a 3-way interaction in the best but not alternative model (Models 2, 23, 25)

| **Model** | **Urbanization Metric** | **Best or Alternative Model** | **Variable** | **AIC** |
| --- | --- | --- | --- | --- |
| 1 | Distance to Urban Center | Best Model | Abundance | 179.929 |
| 2 | Urbanization Score | Best Model | Abundance | 180.808 |
| 3 | Distance to Urban Center | Best Model | Simpson's Diversity | -21.187 |
| 4 | Urbanization Score | Best Model | Simpson's Diversity | -23.829 |
| 5 | Distance to Urban Center | Best Model | Menhinick’s Species Richness | 108.914 |
| 6 | Urbanization Score | Best Model | Menhinick’s Species Richness | 105.806 |
| 7 | Distance to Urban Center | Best Model | Evenness | 24.56 |
| 8 | Urbanization Score | Best Model | Evenness | 23.158 |

**Table S-8** Summary of AIC scores from general linear models examining the effects of urbanization on pollinator community structure (Question 3)

| **Model** | **Urbanization Metric** | **Best or Alternative Model** | **Variable** | **AIC** | **ΔAIC from Best Model** |
| --- | --- | --- | --- | --- | --- |
| 1 | Distance to Urban Center | Best Model | Abundance | 114.907 | - |
| 2 | Distance to Urban Center | Alternative Model | Abundance | 116.709 | 1.802 |
| 3 | Urbanization Score | Best Model | Abundance | 115.009 | - |
| 4 | Urbanization Score | Alternative Model | Abundance | 115.038 | 0.029 |
| 5 | Distance to Urban Center | Best Model | Simpson's Diversity | -28.612 | - |
| 6 | Distance to Urban Center | Alternative Model | Simpson's Diversity | -28 | 0.612 |
| 7 | Urbanization Score | Best Model | Simpson's Diversity | -29.471 | - |
| 8 | Urbanization Score | Alternative Model | Simpson's Diversity | -29.38 | 0.091 |
| 9 | Distance to Urban Center | Best Model | Menhinick’s Species Richness | 72.188 | - |
| 10 | Distance to Urban Center | Alternative Model | Menhinick’s Species Richness | 72.764 | 0.576 |
| 11 | Urbanization Score | Best Model | Menhinick’s Species Richness | 71.547 | - |
| 12 | Urbanization Score | Alternative Model | Menhinick’s Species Richness | 72.384 | 0.837 |
| 13 | Distance to Urban Center | Best Model | Evenness | 12.892 | - |
| 14 | Distance to Urban Center | Alternative Model | Evenness | 13.505 | 0.613 |
| 15 | Urbanization Score | Best Model | Evenness | 12.033 | - |
| 16 | Urbanization Score | Alternative Model | Evenness | 12.124 | 0.091 |

**Table S-9** Summary of AIC scores from general linear models examining the effects of urbanization and proximity to an urban corridor on pollinator community structure (Question 4). All alternative models yielded identical conclusions to the best models

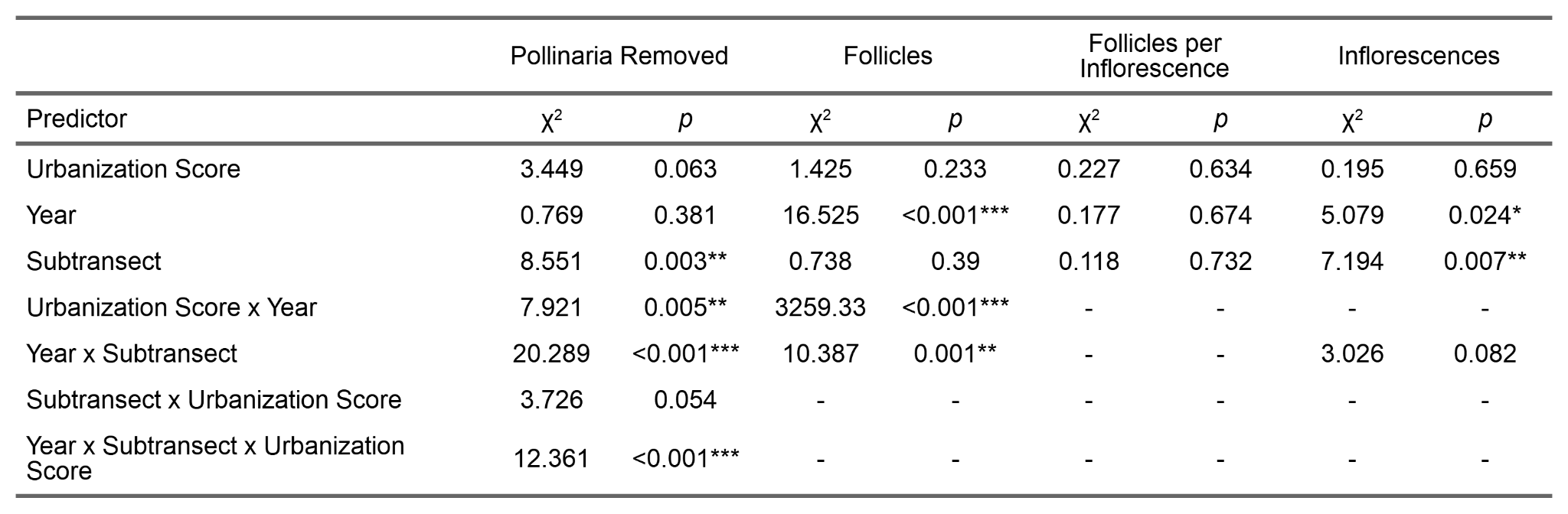

**Table S-10** Results from general and generalized linear mixed models examining the effects of urbanization and proximity to an urban corridor on pollinaria removed, follicles, follicles per inflorescence, and inflorescences. Urbanization was quantified via urbanization score, and only urban populations were included. Shown are maximum likelihood χ^2^ and p-values obtained from a type III sums-of-squares ANOVA
